# A murky ruling threatens the fate of millions of US wetlands

**DOI:** 10.1101/2023.11.14.567035

**Authors:** B. Alexander Simmons, Marcus W. Beck, Kerry Flaherty-Walia, Jessica Lewis, Ed Sherwood

## Abstract

For decades, federal protections were extended to wetlands adjacent to navigable “waters of the US” by the Clean Water Act. In its *Sackett v. EPA* ruling, however, the US Supreme Court redefined the meaning of “adjacent,” eliminating protections to wetlands without a continuous surface connection to these waters (i.e., geographically isolated wetlands). Yet it remains unclear how this continuous surface test will work in reality, where ecological connectivity often extends beyond physical connectivity. Here, we estimate that 22-36% of US wetlands could be considered geographically isolated, depending upon how isolation is defined on the ground, with wetlands in the Midwest and Northeast at greatest risk of losing protection. Stable state-level protections are urgently needed to secure the country’s wetlands from further pollution and destruction.

**One-Sentence Summary:** Over 3 million wetlands could be considered “geographically isolated” and lose extended protections from the Clean Water Act.

## The precarious state of US wetlands

Wetlands perform essential functions within the landscape, such as water storage and filtration, carbon sequestration, and serving as critical habitat, providing hydrological, chemical and biological benefits that sustain ecological and sociological well-being (*1*). In some wetland systems, such as riparian networks and floodplain swamps, the hydrological and ecological connectivity between upstream and downstream waters is obvious. Some wetlands are considered “geographically isolated”–surrounded by uplands without persistent surface water connections–but remain functionally connected to the natural landscape through the movements of plants and animals. These geographically isolated wetlands support terrestrial biodiversity and water cycle dynamics due to their unique hydrological regimes, and serve as important refuges for rare and threatened species (*2*).

The loss of a single isolated wetland can be significant if it supports an endangered species, but the cumulative loss of many of these wetlands can cause regional consequences, reducing landscape connectivity and increasing isolation. In addition to their ecological importance, these isolated wetlands are economically valuable; it is estimated that ephemeral and seasonal-flowing streams provide $15.7 trillion annually in ecosystem services in the conterminous US, and wetlands outside of floodplains contribute an additional $673 billion each year (*1*). For centuries, humans have altered wetland connectivity for flood control, agricultural practices, and development, resulting in a loss of 53% of all wetlands in the conterminous US over a period of 200 years (*3*).

At the federal level, protections for wetlands are secured under the Federal Water Pollution Control Act, also known as the Clean Water Act (CWA), which enables federal agencies to regulate pollutant discharges into “waters of the United States” (WOTUS) and establish surface water quality standards. Historically, most wetlands were considered to fall within the WOTUS definition and thus under protection by the CWA. A wide range of wetland regulations are also present at the state-level, but less than half of all states have explicit protections for geographically isolated wetlands (*1*). For many of the country’s small, isolated wetlands, protections afforded by the CWA have been critical for filling these state regulatory gaps. Over the last decade, however, the WOTUS definition has become increasingly ambiguous, raising concerns over the future of isolated wetlands across the country.

## A murky ruling

After a long history of regulatory pivots throughout the Obama, Trump, and Biden Administrations (*4*), the ruling by the US Supreme Court in the case of *Sackett v. EPA* has managed to halt the swing of the policy pendulum while leaving many questions unanswered. Writing for the majority opinion, Justice Samuel Alito argued that the CWA only applies to wetlands with a “continuous surface connection to bodies that are ‘waters of the United States’” and concluded that geographically isolated wetlands are not covered by the CWA’s extended protections to wetlands that are “adjacent” to waters of the US (*5*). Justice Brett Kavanaugh criticized Justice Alito’s definition for its interpretation of the statutory “adjacent” terminology as effectively “adjoining,” arguing “By narrowing the Act’s coverage of wetlands to only adjoining wetlands, the Court’s new test will leave some long-regulated adjacent wetlands no longer covered by the Clean Water Act” (*6*).

The new ruling defines a wetland under WOTUS jurisdiction as (1) having a continuous surface connection with an existing surface water, and (2) being practically indistinguishable from an ocean, river, stream, or lake where the continuous connection is defined. Many have critiqued this decision with near unanimous agreement that it is not scientifically justified and is legally unprecedented relative to past CWA amendments (*7*). Despite concerns from environmental scientists and legal experts (e.g., *7, 8*), there has not been a comprehensive and quantitative estimate of the amount and extent of wetlands that are now vulnerable to exclusion under the latest ruling. These estimates are critical to understand the potential effects this decision has on the nation’s water quality, in addition to motivating state and local governments to bolster existing protections where federal rulings fall short.

To identify geographically isolated wetlands at risk from the ruling, we obtained spatial data for over 35 million wetlands in the United States mapped by the National Wetland Inventory (NWI) (*9*) and calculated the shortest distance between each wetland and the nearest hydrological feature (i.e. water body or flow line) mapped by the National Hydrography Dataset (*10*). We excluded wetlands from our analysis that are unlikely to be affected by the new WOTUS definition, including (1) wetlands classified as estuarine and marine deepwater habitat, (2) wetlands overlapping an existing protected area with strict mandates for biodiversity conservation (US Geological Survey Gap Analysis Project status 1 or 2)(*11*), and (3) wetlands smaller than 0.25 acres, which would likely remain unprotected or excluded from permitting activities in most states regardless of the WOTUS decision. This resulted in the inclusion of 23.3 million wetlands (61.4% of NWI features) covering 245 million acres (66.4% of NWI area).

The Supreme Court’s recommended “continuous surface connection” test means that any physical separation between WOTUS boundaries and adjacent wetlands could disqualify those wetlands from federal protection. We estimated the number of wetlands that could be considered geographically isolated if 1, 10, 20, 30, 40, 50, 60, 70, 80, 90, or 100 m from the nearest hydrological feature is used as the distance threshold beyond which a wetland would be considered isolated. These ranges provided reasonable bounds on the estimates of amount and extent of isolated wetlands by accounting for both the ambiguity of the SCOTUS definition and uncertainties in the data. Recognizably, natural hydrologic flows are determined through complex interactions between topography, soil infiltration, groundwater influences, and seasonal weather patterns; a straight-line distance does not fully account for these factors, but provides a reasonable approximation in the absence of incorporating additional datasets to more accurately define water flow between hydrologic features. We also obtained data from Creed et al. (*1*) on the presence of regulatory protections for geographically isolated wetlands in each state to better gauge differential risks to wetlands in states with existing, limited, or no protections (Table S1). See Supplementary Material (SM) for additional details on the methodology.

## Wetlands at risk

We estimate that 8.42 million wetlands (36% of all wetlands included) could be considered geographically isolated using the 1 meter distance threshold. This represents nearly 29 million acres, or 12% of the total wetland area evaluated (Fig. 1a). Under a greater threshold (100 m), the number of isolated wetlands could reduce to 5.09 million (22%), covering 15.6 million acres (6%). Protections for the 3.33 million wetlands (14%) within this distance range are the most vulnerable to the ambiguous definitions in the Sackett ruling. Roughly 41% of these wetlands are located in states without existing protections for geographically isolated wetlands, while 15% are in states with some limited protections in place. These isolated wetlands are most prevalent in the northern midwest and northeastern states (Fig. 1b). For example, approximately 50-70% of wetlands in Minnesota and North Dakota could be considered geographically isolated (depending on the distance threshold), representing 24-39% of each state’s wetland area. In the northeast, Maine and Maryland have some of the highest representation of isolated wetlands: 23-49% of their wetlands could be considered isolated, representing 13-26% of their wetland area.

**Fig. 1.**
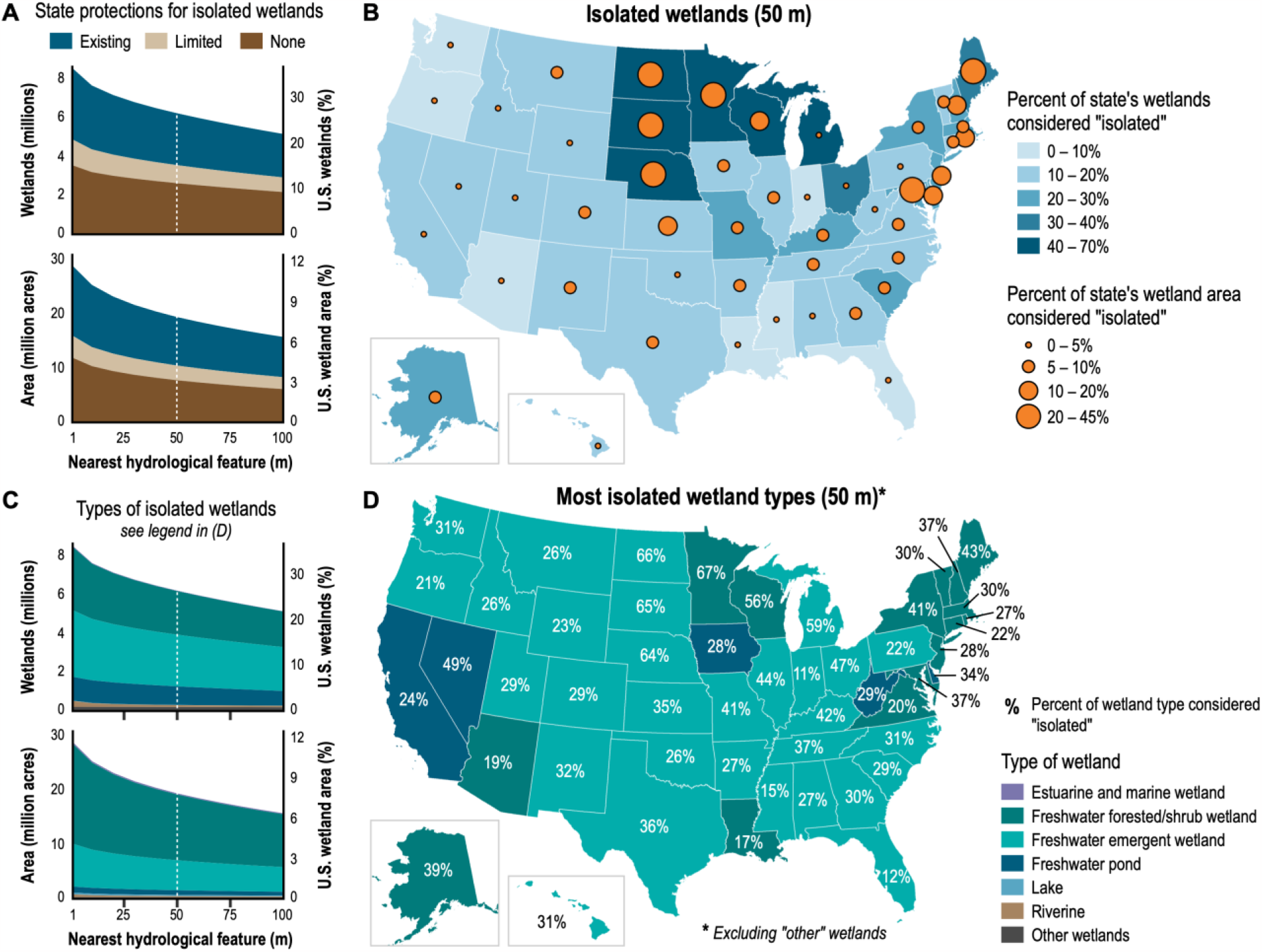
Wetlands at risk under the Supreme Court’s ambiguous definition of wetlands protected by the Clean Water Act. (**A**) The share of US wetlands that could be classified as “geographically isolated” using different definitions of isolation depending upon their proximity to the nearest hydrological feature (1 to 100 m), and distinguished between wetlands in states with existing, limited, or no protections for isolated wetlands. Dotted line indicates the average distance threshold investigated (50 m), which is used in (B) and (D). (**B**) Proportion of each state’s total number of wetlands and total wetland area that could be considered geographically isolated using a 50 m distance threshold. (**C**) The share of geographically isolated wetlands in (A) according to the type of wetland. (**D**) The type of wetland in each state (excluding “other” wetlands) with the greatest proportion considered isolated using a 50 m distance threshold. Percentages show the percent of the respective wetland type that is considered isolated.

The majority of wetlands at risk are freshwater emergent wetlands (e.g. marshes, wet prairies) (Fig. 1c). These wetlands make up 40-44% of all wetlands classified as geographically isolated under the different distance thresholds. Freshwater forested or shrub wetlands (e.g. swamps, hammocks, wet flatwoods), however, have the greatest area at risk: 62-64% of the total isolated wetland area consists of these wetlands. While emergent wetlands are the most frequently isolated across the country, some states deviate from this trend (Fig. 1d). Many northeastern states have a larger share of their forested/shrub wetlands classified as geographically isolated, while freshwater ponds are relatively more isolated in Nevada, West Virginia, Iowa, and California. Notably, 81-93% of “other” wetlands (e.g. farmed wetlands, saline seeps) could be considered geographically isolated—the highest share of any wetland type—but make up less than 1% of all wetlands in the country.

## One protection does not fit all

Environmental policy should be based upon the best available and most broadly accepted scientific standards, definitions, and recommendations. The Sackett ruling affects vital ecological systems and services that the US relies upon for public health, environmental sustainability, and economic success. An impactful federal policy definition, such as “WOTUS,” should at a minimum reference scientifically accepted assessments. The undefined “continuous surface water connection” standard referenced in the SCOTUS majority opinion falls short of providing national to local legal interpretations and policy guidance for ecologically and economically vital wetlands of the US. Although federal guidance states that the NWI and NHD datasets are insufficient for spatial descriptions of WOTUS, primarily due to errors of omission and commission (*12*), this argument is ultimately a red herring. These datasets represent the most comprehensive, national-scale source of information on surface water coverage in the US (*7*), and an objective assessment of wetlands at risk with these data remains highly informative regardless of the data limitations.

Recognizably, the WOTUS definition has changed before and will likely change again. Yet state and local governments should not be reliant upon federal policy to comprehensively protect the varied wetland types represented across the US. State and local governments should take this opportunity to strengthen, expand, or initiate protections for habitats that are threatened by anthropogenic harm and may now fall outside federal protections. However, the opposite approach is now being taken. At the state level, wetland protections are already aligning with this new federal policy. For example, the North Carolina legislature overruled a gubernatorial veto through a supermajority vote in both chambers to repeal existing state-level protections for isolated wetlands just one month after the Sackett opinion was published (*13*). Local protections risk further contraction to this federal standard given continuing economic growth and development pressures.

It is unclear how the Sackett ruling will impact environmental protection regulations in states like Florida, where the Department of Environmental Protection recently took over CWA Section 404 wetland permitting from the Environmental Protection Agency (*14*) despite concerns from the state’s environmental advocacy organizations. Our analyses attempted to categorize risks at the state level, where existing, limited, or no isolated wetland protections could be readily discerned. However, wetland protection standards under more local governance models should not be discounted as important public policy. In the Tampa Bay region of Florida, for example, definitions of “waters of the state and/or county” provide an additional backstop to limiting impacts to isolated freshwater wetlands that provide important migratory bird habitat functions across state jurisdictions, while also being conduits for direct aquifer recharge of regional and state drinking water supplies. Likewise, Maine specifically recognizes and manages unique, isolated wetland types that are identified and valued as rare ecosystems under state policies. Despite some states having more protective regulatory frameworks for isolated wetlands, the Sackett ruling may increase their exposure to future legal challenges, creating additional economic and litigious drains on society.

Scientific consensus remains that wetlands, no matter their size or interconnectivity, are vital to biodiversity and human well-being. While we strive to limit our impacts to these systems through various policies (e.g. “WOTUS” or “no net loss” in the US), globally our efforts are failing (*15*). The Sackett ruling is already leading to cascading policy effects at the state-level, and further erosion of local protections may be on the horizon. Our analyses reinforce that collective protections at the federal, state, and local level will be needed to reverse wetland attrition within the US and to help meet global goals to sustain social and ecological systems.

## Acknowledgments

We are grateful to C. Bohlen, D. Carpenter, and M. Rains for providing insights and perspectives that helped guide conversations during this manuscript’s development.

## Funding

Tampa Bay Estuary Program (TBEP) funding for this work stems from EPA Section 320 Grant Funds, and the TBEP’s local government partners (Hillsborough, Manatee, Pasco, and Pinellas Counties; the Cities of Clearwater, St. Petersburg, and Tampa; Tampa Bay Water; and the Southwest Florida Water Management District) through contributions to the operating budget.

## Author contributions

Conceptualization: BAS

Methodology: MWB, KFW, BAS, ES

Analysis: MWB, BAS

Visualization: BAS

Writing: MWB, KFW, JL, BAS, ES

## Competing interests

Authors declare that they have no competing interests.

## Data and materials availability

All data used for analyses are available from their respective sources. R code for replicating this analysis is available from GitHub (https://github.com/tbep-tech/wetlands-eval). The complete dataset, with files for each state, is hosted on the Knowledge Network for Biocomplexity (https://doi.org/10.5063/F1M043V0).

## Supplementary Materials

Materials and Methods

Table S1

References (*16*–20)

## Notes

https://github.com/tbep-tech/wetlands-eval

https://doi.org/10.5063/F1M043V0

### Competing Interest Statement

The authors have declared no competing interest.

