## Supplementary Material for "A murky ruling threatens the fate of millions of US wetlands"

**The PDF file includes:**

Materials and Methods
Table S1
References *16-20*

### Materials and Methods

Under the new Supreme Court decision, wetlands are jurisdictional to Waters of the United States (WOTUS) if 1) a continuous surface connection is present with an existing WOTUS, and 2) the wetland is practically indistinguishable from an ocean, river, stream, or lake where the continuous surface water connection is identified. A national-scale assessment for the United States was conducted to identify potential geographically isolated wetlands (GIWs), focusing specifically on the first requirement to define those beyond a given straight-line distance from existing water bodies. The second requirement that a wetland must be "practically indistinguishable" cannot be assessed at a national scale with existing datasets. Our analysis further recognized that existing wetland protections at the state level may have precedence over federal protections. All GIWs were considered similarly across states regardless of existing protections, as state rules can now be challenged under the new federal WOTUS definition.

The National Wetland Inventory (NWI) maintained by the US Fish and Wildlife Service and the National Hydrography Dataset (NHD) maintained by the US Geological Survey were used for the assessment. The NWI is the most comprehensive geospatial dataset (1:24,000 scale) of wetlands in the US, representing the combined mapping efforts of states, federal agencies, tribal governments, regional and local governments, and nonprofit organizations. Data are available for over 35 million wetlands classified as "estuarine and marine deepwater," "estuarine and marine wetland," "freshwater emergent wetland," "freshwater forested/shrub wetland," "freshwater pond," "lake," "riverine," and "other." The "other" category includes farmed wetlands, saline seeps, or other miscellaneous types (*16*). The NHD is mapped at 1:24,000 scale and includes line and area features for flow networks and waterbodies, respectively. Both the NWI and NHD are available as separate geodatabases or shapefiles for each state. A custom analysis workflow at this spatial scale was used to quantify the amount and areal coverage of GIWs by state. Both the NWI and NHD are not without accuracy limitations, which primarily include errors of omission/commission based on constraints of the data used to create each layer (*17*, *18*). However, both datasets represent the best estimate of surface water coverage in the US and an assessment of potential GIWs is informative regardless of the limitations.

The analysis was conducted using the open source R statistical programming language (version 4.2.3)(*19*). A custom workflow was developed to iteratively download the NWI and NHD spatial layers for each state to identify GIWs based on the Euclidean distance of wetland features to NHD features. The simple features package, "sf" (*20*), provided the core functions for the spatial analyses, including use of the st_read() function for importing relevant layers from the state geodatabases and calculating distances between features using the st_nearest() function. For the latter analysis, the NWI layer for each state was compared separately to the flowline and waterbody layer from the NHD, resulting in an index of NHD features that were nearest to each NWI feature. The distance between the NWI features and the nearest NHD feature were estimated using the st_distance() function and the minimum distance to either a flowline or waterbody feature was estimated for each wetland. Centroids were also calculated for each wetland using st_centroid() to provide spatial information for each feature and to minimize the storage requirements of the results. The final datasets included tabular information for each state, where each row was an individual wetland, with columns for the wetland attribute, acreage of the wetland, latitude and longitude (WGS 1984) of the centroid, distance of the wetland in meters to the nearest NHD feature, the state abbreviation, and wetland type, available on the Knowledge Network for Biocomplexity at https://doi.org/10.5063/F1M043V0. All geospatial analyses were conducted using the Albers equal area projection with a North American Datum of 1983. Code for the analysis workflow is available on GitHub at https://github.com/tbep-tech/wetlands-eval.

The state tabular data with the distance of each wetland feature to the nearest NHD feature were used to quantify the amount and extent of GIWs using a range of thresholds. In total, 38,011,807 wetlands were assessed, where the distance to NHD features ranged from 0 to 26 kilometers. A range of distances for each NWI feature to the nearest NHD feature were used to develop an expectation of the amount and extent of GIWs at risk in each state. The Supreme Court decision that wetlands must have a "continuous surface connection" to existing navigable waters does not provide specificity or quantitative guidance on how this should be defined. The range of values used to quantify potential GIWs in our analysis varied from 1 to 100 meters (at 10 meter intervals) to take into account the uncertainty in the current definition, as well as the uncertainty in the mapping products used in the analysis. Based on these ranges, the amount and extent of GIWs in each state were quantified, including an assessment of GIWs by wetland type (i.e., freshwater emergent, freshwater forest/shrub, etc.). The most vulnerable wetland type by state was also identified.

Additional wetlands were removed from the analysis if they were considered irrelevant given current legislation or wetland definitions. First, all wetlands identified as estuarine and marine deepwater were removed from the analysis (n= 236,716, or 0.6% of all NWI features). These wetlands are those present entirely in submerged, subtidal environments that are permanently flooded, such as rock bottom, reefs, or other areas that support aquatic life but are not typically considered conventional wetlands based on soil or vegetation classifications. Moreover, these areas typically exist in subtidal habitats of surface waters under existing Clean Water Act protections and are therefore not at risk. Second, all wetlands with surface area less than 0.25 acres were excluded from analysis (n= 12,491,050, or 32.9% of all NWI features). These "small" wetlands represent those that are typically already unprotected and do not require permits for activities that can degrade or eliminate their function. This size threshold varies by state or smaller regulatory jurisdictions. For example, Florida uses a threshold of 0.5 acres (Rule 32-340 F.A.C), whereas Indiana uses a threshold of 0.1 acres (Indiana Code 13-18-22). As such, the 0.25 minimum acreage criterion represents a generic, de minimis threshold that acknowledges most states do not protect small wetlands.

Finally, all wetlands in existing protected areas were not considered in the analysis, such as those in national parks or marine conservation areas. These wetlands were considered not at risk based on the exclusion of activities in protected areas that can degrade or eliminate wetland function. These wetlands were identified by intersection of the wetland centroids with polygons in the Protected Areas Database (PAD-US) maintained by the United States Geological Survey (*11*). This database is a comprehensive inventory of protected areas, including public and voluntarily provided private lands, that are categorized by "GAP status" (Gap Analysis Program) indicating how they are being managed for conservation purposes. Each feature is assigned an integer of 1 to 4 for the GAP status, with decreasing protections for higher numbers. Only polygons with GAP status as 1 or 2 were used to exclude GIWs from the analysis, where those with GAP status 1 have permanent protections and mandated management plans for biodiversity, and those with GAP status 2 are similar but may receive uses or management practices that degrade the quality of natural communities (e.g., suppression of natural disturbances). Conversely, wetlands in GAP status 3 and 4 were not excluded, where protected areas in GAP status 3 may be subject to extractive uses (e.g., logging, mining) and those in GAP status 4 have no mandated biodiversity protections.

**Table S1.**

Level of protection provided to geographically isolated wetlands by state. Adapted from Creed et al. (*1*). ** Reclassified from original source due to legislative changes in 2023.*

| **Existing** | **Limited** | **None** |
|---|---|---|
| California | Illinois | Alaska |
| Colorado | Indiana | Alabama |
| Connecticut | Massachusetts | Arkansas |
| Florida | Michigan | Arizona |
| Hawaii | New Hampshire | Delaware |
| Maryland | Nevada | Georgia |
| Maine | New York | Iowa |
| Minnesota | Texas | Idaho |
| Nebraska | Vermont | Kansas |
| New Jersey | West Virginia | Kentucky |
| New Mexico | | Louisiana |
| Ohio | | Missouri |
| Oregon | | Mississippi |
| Pennsylvania | | Montana |
| Rhode Island | | North Carolina* |
| Tennessee | | North Dakota |
| Virginia | | Oklahoma |
| Washington | | South Carolina |
| Wisconsin | | South Dakota |
| Wyoming | | Utah |
